# Genomic determinants of speciation and spread of the *Mycobacterium tuberculosis* complex

**DOI:** 10.1101/314559

**Authors:** Álvaro Chiner-Oms, Leonor Sánchez-Busó, Jukka Corander, Sebastien Gagneux, Simon Harris, Douglas Young, Fernando González-Candelas, Iñaki Comas

**Affiliations:** Unidad Mixta “Infección y Salud Pública” FISABIO-CSISP/Universidad de Valencia, Instituto de Biología Integrativa de Sistemas-I2SysBio, Valencia, Spain.; Pathogen Genomics, Wellcome Trust Sanger Institute, Cambridge CB10 1SA, UK.; Department of Biostatistics, University of Oslo, 0317 Oslo, Norway.; Helsinki Institute of Information Technology (HIIT), Department of Mathematics and Statistics, University of Helsinki, 00014 Helsinki, Finland.; Swiss Tropical and Public Health Institute, Basel, Switzerland.; University of Basel, Basel, Switzerland.; The Francis Crick Institute, 1 Midland Road, London NW1 1AT, UK.; CIBER en Epidemiología y Salud Pública, Valencia, Spain; Instituto de Biomedicina de Valencia, IBV-CSIC, Valencia, Spain.

**Author notes:** Contributed equally (Álvaro Chiner-Oms, Leonor Sánchez-Busó).

**Keywords:** Bacterial evolution, pathogen genomics, recombination, transmission, two component system

## Abstract

**BACKGROUND:** Models on how bacterial lineages differentiate increase our understanding on early bacterial speciation events and about the genetic loci involved. Here, we analyze the population genomics events leading to the emergence of the tuberculosis pathogen.

**RESULTS:** The emergence is characterized by a combination of recombination events involving core pathogenesis functions and purifying selection on early diverging loci. We identify the *phoR* gene, the sensor kinase of a two-component system involved in virulence, as a key functional player subject to pervasive positive selection after the divergence of the MTBC from its ancestor. Previous evidence showed that *phoR* mutations played a central role in the adaptation of the pathogen to different host species. Now we show that *phoR* have been under selection during the early spread of human tuberculosis, during later expansions and in on-going transmission events.

**CONCLUSIONS:** Our results show that linking pathogen evolution across evolutionary and epidemiological timescales point to past and present virulence determinants.

## BACKGROUND

The increasing availability of population genomics data has allowed an improved understanding of genotypic and ecological differentiation among closely related bacteria[1]. While a species concept *sensu stricto* cannot be applied to bacteria[2] models exists to understand how species can emerge in natural populations. Depending on the evolutionary forces involved, models range from differentiation driven by natural selection and adaptation to different ecological niches (Ecological Species Concept) to differentiation as a result of restricted gene flow that reinforces isolation (Biological Species Concept). In reality, most natural populations show a combination of both processes with certain overlap between habitats (Overlapping habitats model[3]). The study of natural populations and models shows that the emergence of new species is more common among bacterial groups sharing, partially or totally, their habitat, a process also known as sympatric speciation[4]. Processes of bacterial differentiation are often expected to leave measurable genetic signatures in extant genomes including “speciation islands” (regions of high divergence between the nascent species)[3,4]. These genetic signatures hold clues about the key genomic determinants responsible for ecological differentiation of a nascent species from a common genetic pool. However, how these models apply to professional pathogens, particularly those characterized by an obligate association with their host species, has been little explored.

Species of the *Mycobacterium tuberculosis* complex (MTBC) cause devastating morbidity and mortality in humans and animals, which also lead to important economic losses[5]. The MTBC comprises a group of bacteria with genome sequences having an average nucleotide identity greater than 99% and sharing a single clonal ancestor. This includes the predominantly human pathogens referred to as *Mycobacterium tuberculosis* and *Mycobacterium africanum* as well as a series of pathogens isolated from other mammalian species known as *M. bovis, M. pinnipedi, M. antelope, M. microti*, etc. Human-adapted tuberculosis bacilli show a strong geographic association with some lineages being globally distributed (e.g. lineage 4) and others being geographically restricted (e.g., lineage 5, 6, 7)[6,7]. It is assumed that the causes of this variable geographical distribution are both historical (e.g. trade, conquest, globalization) and biological (e.g. interactions with different human genetic backgrounds)[7]. There is limited transmissibility of animal-adapted strains in humans and, conversely, human-adapted strains transmit poorly among animals[8]. Despite the wide range of host species infected by the different members of the MTBC, there is a maximum of ~2,500 single nucleotide polymorphisms (SNPs) separating any two MTBC genomes[9]. The most closely-related bacteria that fall outside of the MTBC include isolates known as *Mycobacterium canettii* (MCAN). MCAN strains differ from MTBC isolates by tens of thousands of SNP, most of them contributed by recombination between strains[10,11]. MCAN strains have been isolated from the Horn of Africa, predominantly from children and often in association with extrapulmonary tuberculosis[12]. Thus, it is assumed that MCAN represents an opportunistic pathogen with an unidentified environmental reservoir[13] opposed to the obligate MTBC pathogen. Genomic comparisons have identified gene-content differences between MTBC, MCAN and other mycobacteria[10,14,15] as well as genetic differences in virulence-related loci[16,17].

Two pieces of evidence suggest that MTBC and MCAN evolved from a common genetic pool in Africa. Strains of the MTBC have an average nucleotide identity (ANI) to MCAN strains of 98% (range 97.71%-99.30%)[10](our own data), suggesting incomplete or recent speciation (an operational species concept identifies 95% as the barrier to delineate species[18]). In addition, most reports suggest lack of on-going recombination between MCAN and MTBC and within the MTBC[19,20] suggesting complete separation (but see [21]). The second piece of evidence comes from phylogeographic and genetic diversity analyses which identified the origin of the tuberculosis bacilli in Africa[6,22], the likely place of origin of MCAN[10,23]. Taken together, the data suggests that ancestral MTBC and MCAN strains at least shared partially the same niche and genetic pool.

Our understanding about the population genomics events mediating the divergence of the ancestor of the MTBC from a common ancestral pool with MCAN is far from complete. The availability of genome sequences from thousands of MTBC clinical strains as well as of close relatives like MCAN enables us to revisit previous evidence about the evolution of MTBC and also to identify molecular signatures of MTBC speciation events as well as new targets for biomedical research.

## RESULTS

We first analyzed the differentiation between MTBC and MCAN by searching for any hallmark of on-going recombination between and within these groups of strains. Previous reports have suggested that there might be limited but significant recombination among MTBC strains[21,24] while others failed to identify measurable recombination events[25]. To revisit this question, we reasoned that including larger available datasets may maximize the signal of recombination signal if exists. We screened a dataset of complete genome sequences of strains from global sources[26](n=1,591). These genomes are representative of the known geographic and genetic diversity of the MTBC (Additional file 1: Figure S1). Among those genomes, we identified all the variant positions and, more specifically, potential homoplastic sites, i.e. polymorphic sites showing signs of convergent evolution. A total of 96,143 variant positions were called in the 1,591 strains. Parsimony and likelihood phylogenetic mapping of those positions revealed 3,439 positions as being homoplastic (3,6%). Homoplasy can arise as a consequence of recombination but it may be caused by other processes, such as positive selection, sequence gaps contributing to homoplastic counts or mapping/calling errors. To increase the likelihood for homoplastic positions to be due to recombination events, we filtered out non-biallelic positions (1.8%) and potential mapping errors identified by generating synthetic reads around each SNP position (0.2%). In addition, known drug-resistance positions were filtered out due as these are well known instances of convergent evolution[27–29](see Methods).

As a result, a total of 2,360 core homoplastic sites were identified across the 1,591 strains analyzed (2.5% of all variable sites). Homoplastic sites were not significantly accumulating in any region of the genome, suggesting the absence of recombination hotspots (Figure 1A). In addition, with these variant positions, we looked at consecutive runs of homoplastic sites in the genomes. The 2,360 homoplastic positions defined 97 homoplastic runs (see Methods for definition of a homoplastic run, Additional file 1: Figure S2). If the accumulation of homoplasies was due to recombination, we expect the variant positions involved in each consecutive run to share the same phylogenetic history. From the 97 homoplastic runs, we detected only 2 cases in which two variant positions shared phylogenetic congruence. The two regions accounted for 4 convergent variants (Additional file 2: Table S1) and affected strains from different MTBC lineages. Variants in positions 2,195,896 and 2,195,899 fell in the primary regulatory region of *mazE5*[30]. Variants in positions 2,641,161 and 2,641,163 fell in the intergenic region of *glyS* and Rv2358. Although we cannot completely discard the possibility that these represent recombination events, it is more likely that the two regions have been under positive selection, a mechanism known to lead to the accumulation of homoplastic accumulation in the MTBC[29]. In summary, this large-scale variant-by-variant analysis was not able to identify significant on-going recombination between any of the 1,591 MTBC strains analyzed.

**Figure 1.**
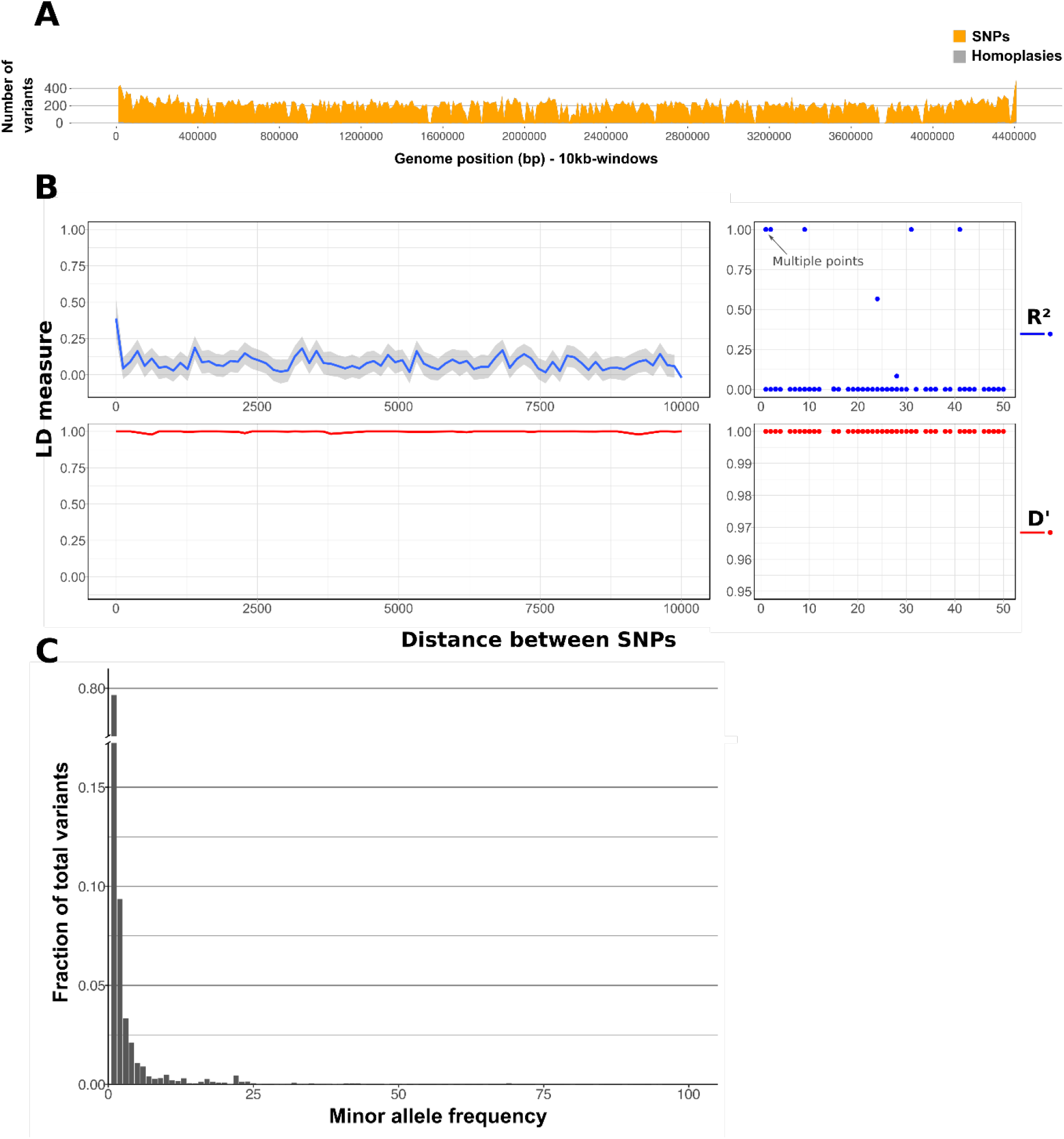
No ongoing recombination within the MTBC. A) Number of homoplasies (grey) as a function of the total number of variants detected (orange) in the MTBC dataset (n=1,591) B) Linkage disequilibrium as a function of genetic distance detected in a representative sample of *Mycobacterium tuberculosis* complex strains (n = 1,591). C) Site frequency spectrum of MTBC strains using the core variant positions.

Due to the low diversity within the MTBC, we also followed alternative approaches to try to identify recombination events with a high statistical confidence. Using an additional method, we evaluated linkage disequilibrium (LD) as a function of the distance between the 94,780 core variant positions. R^2^ has been used in a previous publication to show on-going recombination at very short distances (less than 50 bp.)[21]. In our much larger dataset, R^2^ values were also slightly higher at shorter distances, which is compatible with recombination involving larger fragment sizes. However, a close examination reveals that the peak at short distances is misleading, as it is driven by only six points out of more than 11,000 comparisons (Figure 1B). In addition, R^2^ values are known to fail to reach the theoretical maximum of 1 when variants compared have very different frequencies[31]. This is likely the case for MTBC, in which there is a strong skew of the site frequency spectra towards low frequency values (Figure 1C)[32]. Thus, as an alternative we calculated D’. In this dataset, as expected for a mostly clonal organism, LD measured by D’ remained at its maximum value, even when focusing on distant variant positions more than 5 Kb apart, suggesting very little or no ongoing recombination (Figure 1B).

To further validate these findings, we ran Gubbins with the same dataset and validated them with RDP4 (see methods for details). Gubbins detects the accumulation of a higher than expected number of variants in addition to homoplastic sites as a hallmark of possible recombination. We partitioned the 1,591 strain dataset into the different lineages and screened for possible tracks of recombination. Gubbins reported potential recombining regions characterized by a higher than average number of SNPs however the RDP4 methods did not confirmed any of them. Thus those events maybe real but cannot be confirmed by alternative approaches.

Having established that recombination has little impact on the overall MTBC genetic diversity, we compared a representative dataset of MTBC genomes[6] (n = 219) with 7 MCAN genomes to identify and quantify eventual ongoing recombination within MCAN and between MCAN and the MTBC. Of the 93,922 polymorphic sites identified, 22,718 were biallelic homoplasies (24.2%). The genomic distribution of variant positions and homoplasies in the MCAN group show a landscape different to the MTBC group (Figure 2B). A total of 22,464 (98.9%) of those homoplasies were found only among MCAN strains, representing almost half of the variability within this group (22,464/52,392 biallelic sites, 42.9%) which points to recombination as a main source of variability in MCAN. This is consistent with previous reports[10]. This profile is in sharp contrast with the flat homoplastic profile for the MTBC described above (Figure 1A).

**Figure 2.**
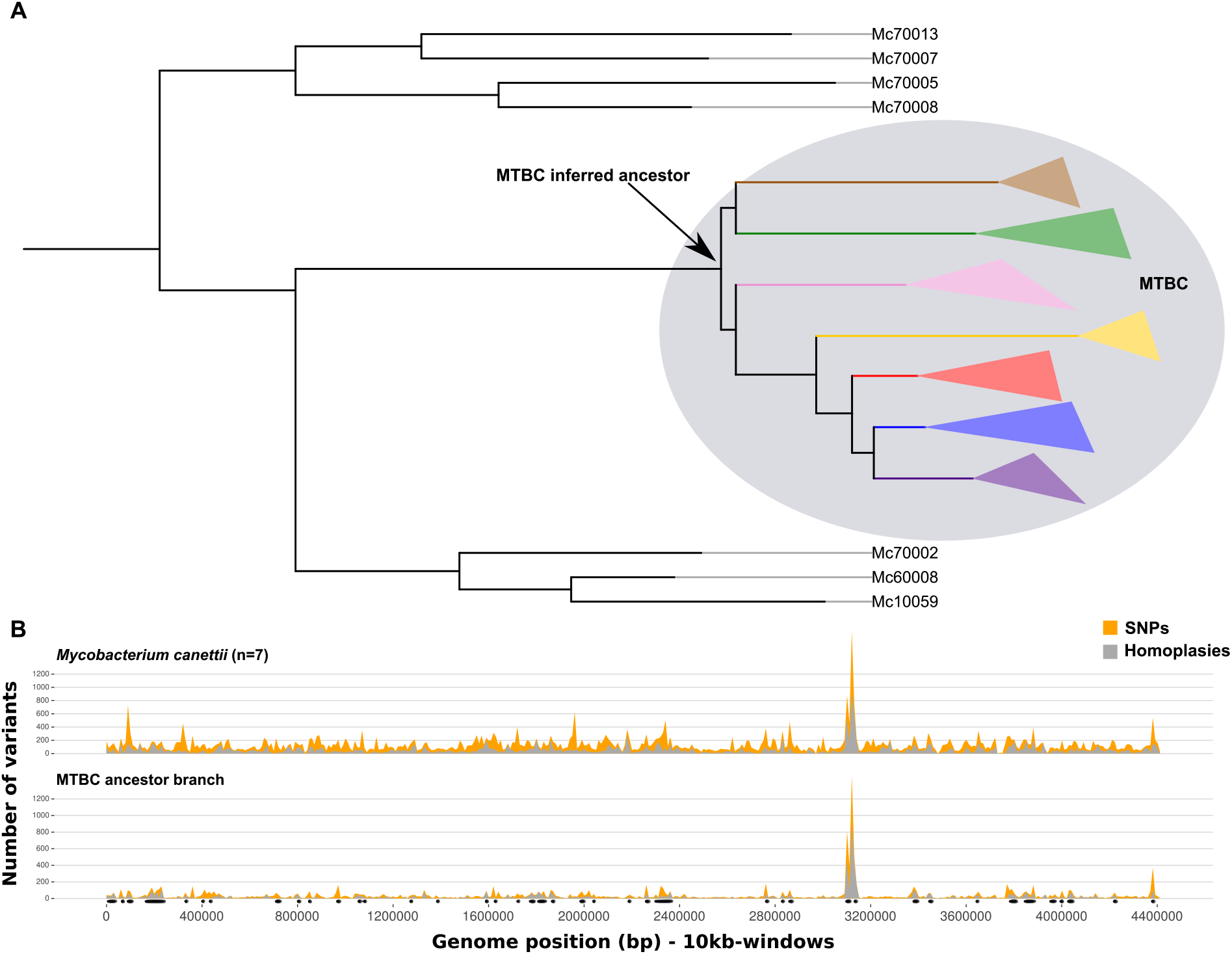
Genome-wide variant profiles vary between *M. canettii, M. tuberculosis* and the MTBC ancestor. A) Schematic view of the phylogenetic relationships between the MCAN groups and the MTBC. In Additional file 1: Figure S3 a maximum-likelihood phylogeny of the MCAN group including the MTBC ancestor can be found. B) Number of homoplasies (grey) as a function of the total number of variants detected (orange) in the MCAN dataset and in the branch leading to the MTBC most recent common ancestor. Black dots indicate recombination events detected in the branch leading to the most recent common ancestor of the MTBC.

To test for ongoing recombination between MCAN and extant MTBC, we identified homoplasies involving both groups. From the 93,922 total variants, 7,934 involved MCAN and MTBC strains. We found 234 biallelic homoplasies involving extant MTBC and MCAN strains, thus compatible with on-going recombination but also with independent diversification. The vast majority of homoplasies detected (97%) mapped to the branch leading to the MTBC clade (thus fixed within the MTBC but variable within the MCAN group). These results indicate that measurable recombination events were common between MCAN and the ancestral branch of the MTBC, but are unlikely during subsequent diversification of the MTBC even when the sample size is greater than previous reports.

### Sympatric and stepwise emergence of the MTBC ancestor

Our results show that recombination with closely related mycobacteria occurred during the emergence of the common ancestor of the MTBC. To gain a better insight on how it occurred we reasoned that instead of comparing MCAN strains against extant MTBC strain we should compare against a reconstructed most common ancestor of the MTBC (derived in Comas et al., 2010[33]). This strategy allowed us to focus on those changes specifically happening in the ancestral branch of the MTBC (see Figure 2A and Additional file1: Figure S3, Figure S4). As described by others, the phylogeny suggests a specific clone of the MCAN group diverged and resulted in the MTBC[11,23]. To do so we extracted all the variant positions that were homoplastic between the MTBC ancestor and any of the MCAN strains (7,700 positions). The SNPs mapping to the branch leading to the MTBC ancestor genome showed a similar homoplasy profile to that of the MCAN strains (Figure 2B), suggesting that there were not hard barriers to gene flow between ancestral MTBC and MCAN ancestral strains, thus supporting a model of sympatric speciation. Notably, both MCAN and the MTBC ancestor shared a peak around the CRISPR region, highlighting the dynamic nature of this region possibly as a result of common phage infections.

A Gubbins analysis including MCAN genomes and the most likely common ancestor of the MTBC revealed a total of 65 recombination events mapping to the branch leading to the MTBC (Additional file 2: Table S2). Mapping of variants into the phylogeny revealed that those regions were coincident with a high number of homoplastic variants between MCAN and the MTBC (Additional file 1: Figure S4). To explore whether these fragments reflected real recombination, we constructed a phylogeny for each of them. A comparison with the topology of the non-recombinant alignment (whole genome alignment subtracting the recombinant regions) using those recombinant regions with enough phylogenetic signal (Additional file 1: Figure S5) revealed significant incongruence (SH test; p-value<0.05, Additional file 1: Figure S6, Additional file 2: Table S3, Additional file 1: Supplementary text). Thus, both Gubbins and phylogenetic approaches indicated that these 65 regions are likely recombinant regions.

The potential recombinant regions were dated using BEAST and the results showed that their inferred ages differ from the established time since the Most Recent Common Ancestor (tMRCA) of the MTBC (see Additional file 1: Figure S7 and Additional file 1: Supplementary text for details). Although the distribution of tMRCA for the fragments represents a continuum, the analysis suggests a separation between “recent” recombination events and “ancient” events closer to the time of divergence from the MCAN group (Figure 3A). The large HPD intervals preclude any firm conclusion but the results suggest that some regions in the MTBC ancestral genome were restricted to gene flow before than others (Additional file 1: Figure S7).

**Figure 3.**
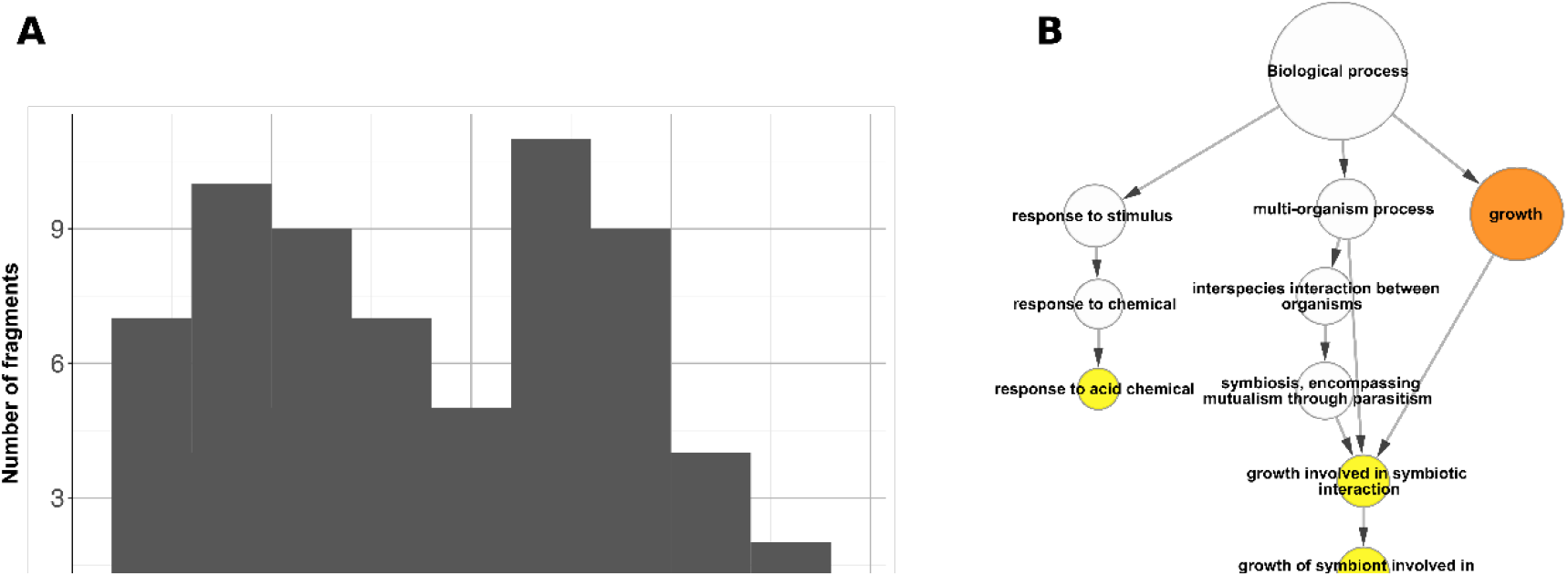
Past recombination between *M. canettii* strains and the MTBC ancestor. A) Histogram distribution of the recombination fragments relative ages. The age was relativized from 0 to 1, being 0 the age of the non-recombinant fraction of the genome and 1 the highest value in the confidence intervals (Figure S5). B) Gene Ontology terms overrepresented in the coding regions contained in the recombinant fragments.

If recombination played a major role in shaping the MTBC ancestral genome with regards to pathogenesis, we would expect some functions related to the interaction with the host to be affected. Indeed, we observed an enrichment in experimentally confirmed essential genes in the regions involved in recombination events, suggesting that recombination targeted important cell functions (Chi-square test; p-value < 0.01). An enrichment analysis of Gene Ontology terms for the genes contained in these regions identified functions related with growth, and most specifically with the category “growth involved in symbiotic interactions inside a host cell” as significantly overrepresented (Binomial test; adj. p-value < 0,05) (Figure 3B). This category can be interpreted as genes involved in a strong association between the pathogen and the host. Remarkably most of the genes involved have been implicated in virulence using animal models of infection (see Discussion, Additional file 2: Table S4).

The recombination profile shown in Figure 2 suggests that the MTBC ancestor recombined with MCAN ancestral strains and, thus, they shared a common niche. A sympatric model of speciation predicts that some parts of the genome will be involved in adaptation to a new niche[4]. The hallmark trace would be the accumulation of variants differentiating the emerging species, at the genome-wide level or in a few loci, as a consequence of reduced recombination between both groups. We identified all the variants that mapped to the ancestral branch of the MTBC and that had a different nucleotide in all the MCAN strains, the so called divergent variants (divSNPs). The distribution of divSNPs per gene revealed that only few of them accumulated divSNPs in the branch leading to the ancestor while most genes did not (Figure 4A). This pattern is compatible with population differentiation models in which niche overlap between emerging species is high[3]. The genome-wide landscape of divergent variants (n = 5,688, Figure 4B) revealed that a total of 120 genes harbored more divergent variants than expected by chance (see Methods)(Figure 4B).

**Figure 4.**
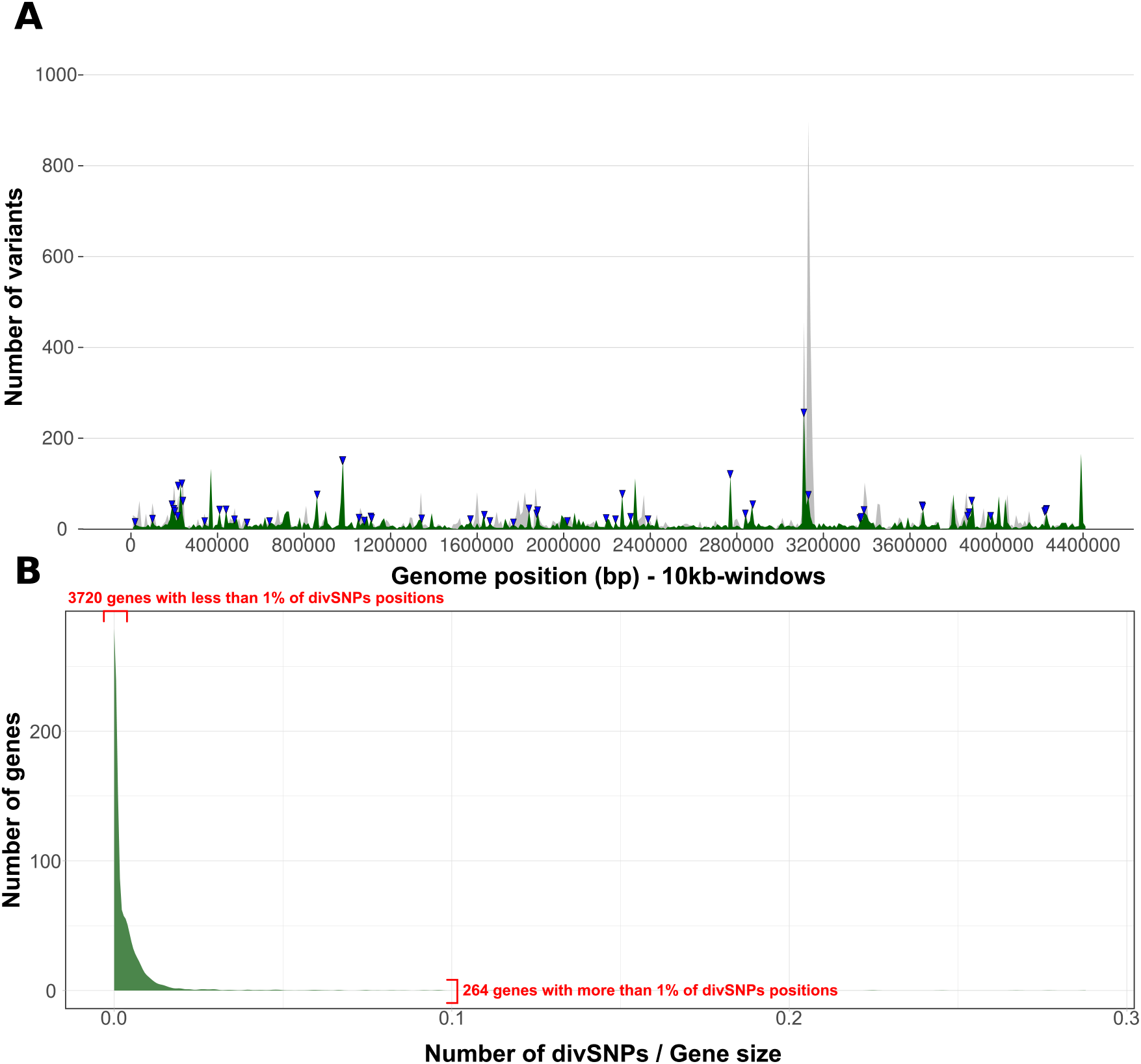
Divergent positions between the MTBC ancestor and *M. canettii* clade. A) Average of divSNPs per 10 kb positions (green) as compared to the average of homoplastic variants (gray). Blue arrows above the distribution are genes that significantly accumulate more divSNPs. B) Accumulation of divSNPs per gene, corrected by gene length. A small number of genes accumulate a high amount of divSNPs while most of the genes have a low number of variants or even none. This pattern resembles those of high habitat overlap derived from Overlapping Habitat Models(3).

However, bacterial genomes are highly dynamic and different processes can contribute to the genetic make-up of extant species. Consequently, not all the detected regions necessarily result from pure divergence by accumulation of substitutions. A detailed comparative genomics and phylogenetic analysis using available genome databases of bacteria identified several genes in which divSNPs were introduced by horizontal gene transfer (n = 12) or by recombination to a MCAN not present in our dataset (n = 54) and one with uncertain phylogenetic origin (see Additional file 1: Supplementary Text for detailed results). Thus, a total of 53 genes in the MTBC ancestral genome were highly divergent with respect to MCAN due to substitution events (Additional file 2: Table S5). While the genome-wide analysis identified divSNPs that might result from genetic drift or hitch-hiking events associated with selection on other loci, their accumulation in only 53 genes suggests that those regions might have played an important role during the process of niche differentiation. In agreement, those 53 genes are significantly more conserved than the rest of the genome (dN/dS = 0.154 vs genome average dN/dS = 0.279, chi-squared p-value <= 0.001). This result suggests that, despite the increased divergence from the MCAN strains, those 53 regions have been evolving under purifying selection. Alternatively, the accumulation of divergent variants could also represent hotspot regions for mutation. None of the genes showed a similar pattern of mutation accumulation in other MCAN (no overlap between the divSNPs probabilities distributions for these 53 genes and the rest of the genomes, t-test p-value < 0.05).

### Regions under positive selection after the transition to obligate pathogen

Having established that some divSNPs accumulate in genes under purifying selection, we screened for positive selection patterns to identify additional genes relevant in the transition from a newly emerged pathogen to a globally established pathogen. We first revisited the evolution of antigenic proteins. Those regions are recognized by the immune system and most of them are hyperconserved within the MTBC[33,34]. Interestingly, and in agreement with previous data from MCAN genomic analyses[10], the dN/dS calculated in the branch leading to the ancestor showed a very similar pattern, with essential genes being more conserved than non-essential ones and T-cell epitopes being hyperconserved (Additional file 1: Figure S8). Only nine divSNPs (5 synonymous and 4 non-synonymous) were found in T-cell epitope regions, which is significantly less than expected by chance (Poisson distribution, p-value < 0.001).

Thus, antigenic regions do not show an altered pattern or intensity of selective pressure. We then explored what other regions of the genome changed significantly in selective pressure by comparing the MTBC ancestor dN/dS and the actual dN/dS in extant populations using our global reference dataset of 4,598 MTBC strains. We calculated a dN/dS for all the genes with at least one synonymous and one non-synonymous mutation for each of the two sets (divSNPs versus within MTBC SNPs). Due to the low number of divSNPs in individual genes, only 499 genes were evaluated. Consequently, although additional genes to those shown in the ensuing analyses may have changed the selection pattern or intensity, they cannot be evaluated properly (Additional file 2: Table S6). We were particularly interested in those genes with a drastic change from purifying (dN/dS < 1) to diversifying or positive selection (dN/dS >1) or vice versa.

Most of the genes evaluated did not show any sign of changing selective pressure or pattern. However, when looking at the dN/dS variation data, 14 genes appeared as outliers (as defined by Tukey’s method[35])(Figure 5A). Genes Rv1244 (*lpqZ*), Rv3910, Rv0166 (*fadD5*), Rv0874c, Rv1152, Rv1678, Rv1951c, Rv2584c (*apt*), Rv3026c, Rv3276c (*purK*), Rv3370, Rv3759c (*proX*) and Rv3900c were under a stronger negative selective pressure following speciation. Many of them are annotated[36] as hypothetical conserved proteins. On the other hand, only one gene changed to evolve under positive selection after divergence from the MTBC ancestor Rv0758, also known as *phoR*. Notably, PhoR forms part of the PhoP/PhoR virulence regulation system[37]. In the branch leading to the MTBC ancestor, this gene was as conserved at the amino acid level, as other essential genes (Chi-square test; p-value 0.4721), but when we looked within the extant MTBC diversity, the gene was significantly less conserved at the amino acid level than essential genes (Chi-square test; p-value < 0.001).

**Figure 5.**
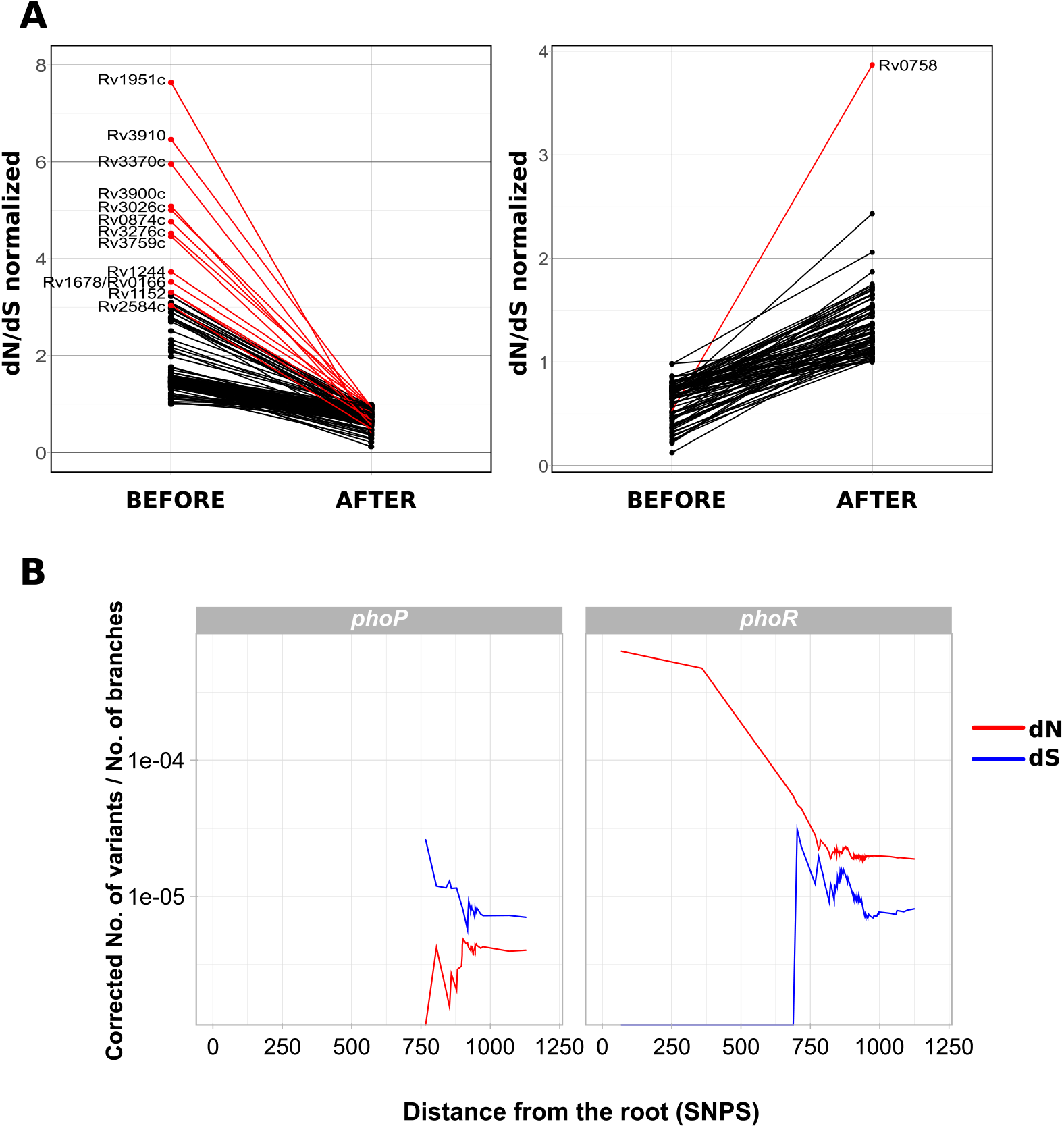
Genes with differential selective pressures across the MTBC speciation stages. A) Genes changing selective pressure in the branch of the MTBC ancestor as compared to extant MTBC strains. Red lines mark those genes being outliers of the dN/dS variation distribution. B) *phoR* and *phoP* show different selective pressure dynamycs. In both cases the accumulation of nonsynonmous (dN) or synonymous (dS) mutations through time is measure as the distance to the most common ancestor of the MTBC. The dN and dS values have been corrected by the number of branches in the phylogeny at each time point.

### Positive selection on *phoR* linked to on-going selective pressures

Given the known central role of PhoPR in MTBC virulence, we focused our attention on the new mutations found in *phoR*. Previous work identified a basal phylogenetic mutation in *phoR* experimentally linked to animal host preferences. Here we report 193 nonsynonymous mutations and 31 synonymous mutations exclusive of human 351 isolates and mapping to very different phylogenetic depths (Figure 6A). The average dN/dS for this gene was well above 1 (dN/dS = 2.37), suggesting the action of positive selection. Furthermore, a plot of the dN and dS values over time reveals that the overall dN/dS remained high along the evolutionary history of the MTBC (Figure 5B) in comparison with the DNA binding response regulator (*phoP*), corroborating that *phoR* has been likely under pervasive positive selection. Codon-based maximum likelihood tests of positive selection are normally powerless in intraspecies comparisons however for the case of *phoR* the tests identified a higher dN/dS than expected by chance and at least two codons with strong evidence to be under positive selection (Additional file 2:Table S7). Additional evidence for the action of positive selection on this gene derives from nonsynonymous mutations, among which we found 34 homoplastic variants, which are strong predictors of positive selection in MTBC (Additional file 2: Table S8). Nonsynonymous mutations significantly accumulated in the sensor domain (Chi-square test, p-value < 0.01), further supporting the hypothesis that they could be involved in the fine-tuning of the PhoR sensitive function to the changing environment during infection (Figure 6C).

**Figure 6.**
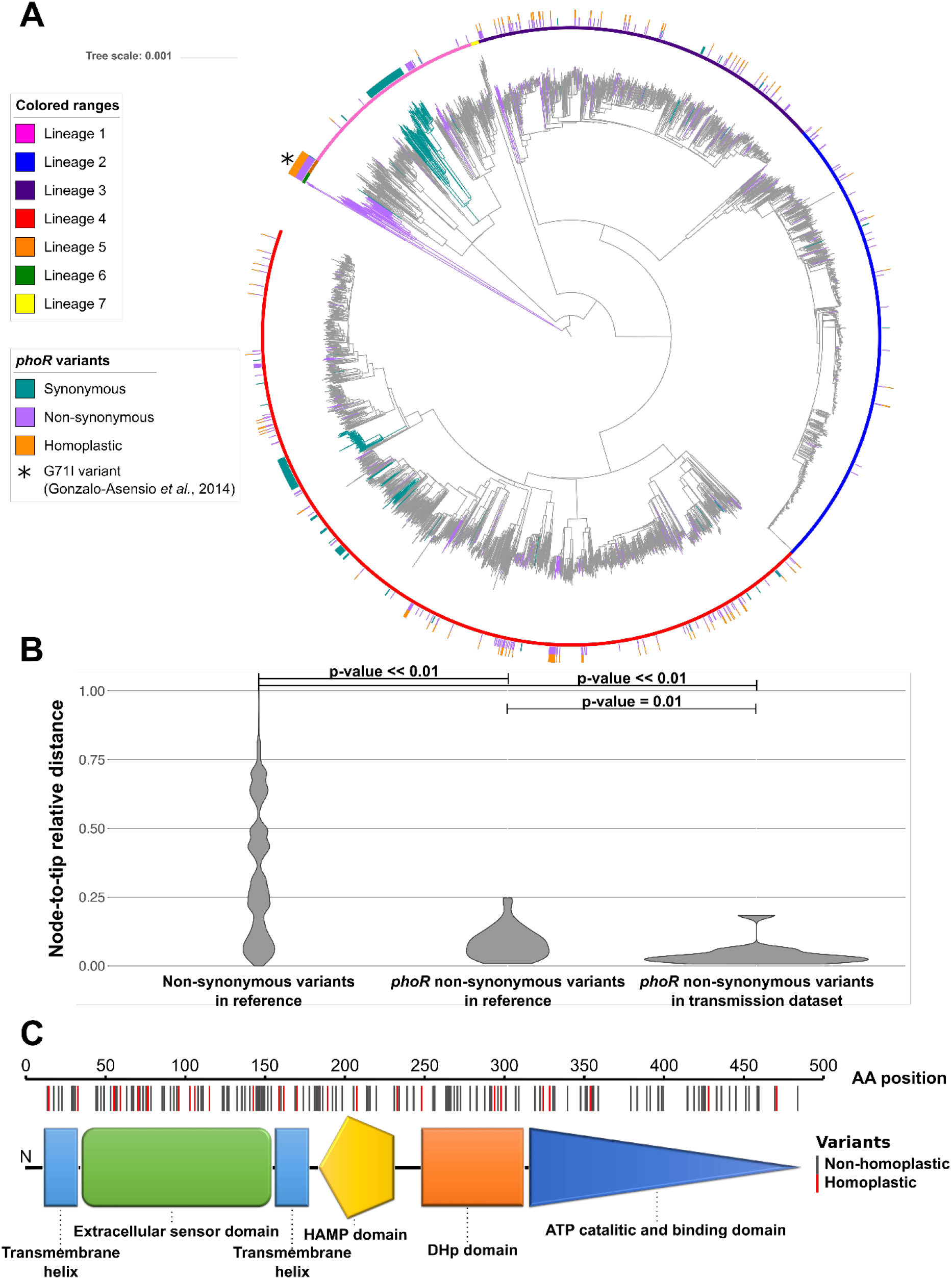
*phoR* is under positive selection in human affecting strains. A) Genome-based phylogeny calculated from a total of 4,598 clinical samples obtained from different sources. The synonymous and nonsynonymous variants found in *phoR* are mapped to the corresponding branch. Variants in internal branches affect complete clades which are colored in the phylogeny. Homoplasies are marked in the outer circle of the phylogeny. The star marks the G71I *phoR* variant common to lineages 5 and 6 previously reported by Gonzalo-Asensio et al.(38) B) Relative ages distribution of the *phoR* variants in the reference dataset from Coll et al.(25) and the transmission dataset(39) in comparison with the rest of the genome variants. C) Schematic view of PhoR with the amino acid changes found across the 4,598 samples dataset marked on it. Amino acid changes are significantly more abundant in the sensor domain (p-value < 0,01).

All the new mutations identified in our analysis were found in human clinical isolates and mapped to relatively recent branches in the MTBC phylogeny (Figure 6A). Thus, we reasoned that most mutations were associated with recent selective pressures as opposed to the previously reported mutations found in *Mycobacterium africanum* lineages 5 and 6, and the animal clade[38] that map to deep branches in the phylogeny (Figure 6A). To prove it, we tested whether novel *phoR* mutations are also arising in clinical settings during infection and recent transmission events. We used a population-based dataset from Malawi[39] where more than 70% of the strains were collected during fifteen years and genome sequenced (n = 1,187). We found 14 mutations (13 nonsynonymous and 1 synonymous) in *phoR* exclusive of the Malawi dataset with *phoR* having a dN/dS of 3.93. Moreover, the mean relative age of the nonsynonymous *phoR* variants was significantly younger than that of other nonsynonymous variants in the dataset (Welch’s t-test, p-value < 0.01) and the *phoR* variants from the Malawi dataset were more recent than those *phoR* mutations from the reference dataset (Welch’s t-test, p-value = 0.01)(Figure 6B). From the 13 nonsynonymous mutations in the Malawi dataset, 8 were markers of recent transmission clusters. Moreover, *phoR* mutations in the Malawi dataset involved larger transmission clusters than other mutations (permutations test, p-value < 0.001). Taken together, these data indicate that novel *phoR* mutations arise during infection and propagate during on-going human-to-human transmissions in clinical settings.

## DISCUSSION

We present evidence that the MTBC ancestor transitioned to an obligate pathogenic lifestyle in sympatry from a common genetic pool including the ancestors of extant MCAN strains. It was already shown before the high recombination rate in MCAN compared to MTBC [10,11]. However, our analyses is different as it specifically focus on the branch leading to the MTBC by comparing MCAN to a reconstructed MTBC common ancestor. Specifically, we found common patterns of genome-wide recombination in the branch leading to the MTBC ancestor and the extant MCAN strains. The high recombination rate between MCAN strains, including the MTBC ancestor, stands in sharp contrast to the strictly clonal population structure of extant MTBC strains. By analyzing events leading to the transition from a recombinogenic to a clonal organism, we have also been able to identify genomic regions under different selective pressures. The comparison between selective pressures before and after becoming an obligate pathogen also allow us to propose PhoR as an important player in the past evolutionary history of the MTBC as well as in current clinical settings.

Population genomics data has led to the development and testing of different models of how different genetic clusters of the same species can arise in sympatry[1,3,40]. In the case of *Vibrio cholerae*, an appropriate combination of certain virulence-associated variants, ecological opportunity and additional virulence factors mediated the successful transition of a particular clone from an environmental to a pathogenic lifestyle[41]. Other known cases such as pathogenic *Salmonella*[42] or *Yersinia* species[43] may have followed a similar trajectory. The MTBC represents an extreme case of clonal emergence associated to its obligate pathogenic lifestyle. Here, we have shown that, despite the high average nucleotide identity between MCAN and the MTBC, there is complete genomic isolation between these organisms. There is experimental evidence that genetic exchange among MCAN strains occurs easily but not between MCAN and the MTBC[17]. We have shown that there is no measurable on-going recombination among the MTBC strains based on our analysis of 1,591 genomes and which is in agreement with other recent reports[19,44]. It is important to note that due to the low divergence within the MTBC, most methods to detect recombination are limited. Hence, we cannot completely exclude the possibility that we might have missed some recombination events. It was previously suggested that recombination (or gene conversion) could be affecting PE/PPE genes disproportionally[45]. Unfortunately, short reads cannot be properly mapped to those regions so our approach does not allow to test this possibility. However, if recombination does occur in the MTBC, it seems to have a minor impact on the overall genetic diversity of the MTBC. Recombination in natural populations depends both on the capacity of chromosomal DNA exchange between the two groups involved and on the ecological opportunity. The mechanisms, if any, by which the MTBC bacilli lost their capacity to recombine while the ancestral genetic pool showed very similar recombination patterns to MCAN strains, remains to be elucidated. Ecological opportunity may also influence on the lack of opportunities of exchange between MTBC strains. Despite the occurrence of super-infections, the bacilli occupy mainly an intracellular lifestyle, thereby reducing the opportunities for genetic exchange.

We can only speculate on how the transition from a likely environmental or opportunistic pathogen to an obligate pathogen occurred, but our analysis has identified a series of non-random evolutionary events. Notably, those events involve core pathogenesis genes. We have identified highly divergent regions in the MTBC ancestor compared to MCAN. The pattern of SNP accumulation suggests that those regions were important in the transition to a closer association with the host. In addition, recombination events mapping to the branch leading to the MTBC ancestor affected essential genes as well as genic regions known to be involved in host-pathogen interaction. The *mymA* operon (Rv3083-Rv3089) is related to the production of mycolic acids and its disruption leads to an aberrant cell-wall structure. Importantly, knock-out studies[46] have shown that this operon is essential for growth in macrophages and the spleen of infected mice. Furthermore, the deletion of genes in this operon leads to a higher TNF-alpha production, highlighting their role on regulating host-pathogen interactions[47]. The other major operon identified in our analysis is the *mce1* operon[48]. *mce1* knock-out mutants are hypervirulent in a mouse model of infection and lose the capacity of a proper pro-inflammatory cytokine production that is needed for the establishment of the infection[49] and granuloma[48]. How these processes are mediated by *mce1* is still not clear pointing to *mce1* as a priority target for biomedical research.

Our analysis identified one gene, *phoR*, which is under positive selection in extant MTBC strains although it was under purifying selection in the MTBC ancestor. PhoR is the sensor component of the PhoPR two-component system, which plays a major role in MTBC pathogenesis[50]. Previous experimental data show that 1) PhoPR is a major virulence determinant in MTBC[37]; 2) that deep phylogenetic branching mutations in PhoPR were involved in the adaptation of the pathogen to different mammalian hosts[38] and that there is at least one case in which natural overexpression of PhoPR in a *Mycobacterium bovis* clinical isolate was linked to a highly transmissible and virulent phenotype in humans[50]. In fact, mutations affecting the whole animal clade in *phoR* have been proposed to be related with the fine-tune MTBC virulence across different animal host species. We find some of these amino acid changes experimentally tested by Gonzalo-Asensio et al. (2014) to have been selected multiple times in unrelated human isolates. Based on these findings, we speculate that recent *phoR* mutations help to fine-tune the immunogenicity of the pathogen during infection, allowing it to manipulate the human host responses and increase the chances of transmission. However, we still need to understand the stimuli and the molecular pathways that are at the basis of the selective pressure driving the evolution of *phoR*. Given that PhoPR is involved in membrane composition[51], mutations in this regulator might also be involved in the susceptibility to some antibiotics. However, antibiotic selection is an unlikely explanation for the oldest mutations in PhoPR, as they likely predate antibiotic usage.

Based on our findings, a model can be proposed in which recombination, together with the acquisition of new genetic material[14,52], generated a favorable genetic background for the MTBC ancestor to occupy or increase its association with mammalian hosts. We see this emergence only once in the MTBC, perhaps because the right combination of multiple, fortuitous genetic events and the particular ecological conditions has occurred only once. More provocative is the idea that MTBC might just be part of a spectrum of association to the host occupied by the different MCAN-MTBC groups. The fact that the so-called Clone A of MCAN strains are more common in the clinic may suggest differences in ecological niches within the MCAN group[23]. In agreement, previous publications[10,23] and our own analysis (Additional file 1: Figure S4) have identified Clone A strains as the closest MCAN evolutionary group to MTBC.

## CONCLUSIONS

In the MTBC, the strong and obligate association with new host(s) was accompanied by new selective pressures. In accordance, we identified genes in the MTBC genome highly diverging from MCAN and evolving under purifying selection, suggesting that they have become essential following MTBC’s transition to an obligate pathogenic life-style. In the final stages of adaptation, positive selection on genes such as *phoR* and others[53–55] likely led to a narrowing of the host-range and later still to a further fine-tuning during the spread of the bacteria within the new host species.

## MATERIALS AND METHODS

### Datasets used

#### *Mycobacterium canettii* dataset

The *M. canettii* dataset is composed by seven draft genomes downloaded from GenBank (CIPT 140010059, NC_015848.1; CIPT 140060008, NC_019950.1; CIPT 140070008, NC_019965.1; CIPT 140070002, NZ_CAOL00000000.1; CIPT 140070005, NZ_CAOM00000000.1; CIPT 140070013, NZ_CAON00000000.1 and CIPT 140070007, NZ_CAOO00000000.1).

#### MTBC global dataset

This dataset includes the global representatives published by Coll *et al*.[26] (n= 1,598), a transmission dataset from Malawi (see below), and the dataset from Walker *et al*.[56] with additional sequences from Europe. The total number of sequences in this dataset when putting together was 7,977 genomes. As the datasets from Walker *et al*. 2015 and Guerra-Assunção *et al*. 2015 are enriched in epidemiological clusters, we identified all clusters at a maximum distance of 15 SNPs (common threshold in MTB epidemiology), removed samples potentially coinfected with more than one strain and then kept just one representative from each cluster. Thus, the final number of genomes for this dataset was 4,598. Out of this total, we used different subsets of strains depending on the analysis to be performed. A 1,591 sequences subset (corresponding to Coll *et al*. 2014) was used for the recombination analyses within the MTBC. A specific subset of 219 sequences, corresponding also to global representatives, was used for Gubbins because it was not computationally feasible to run the program with more strains. The total 4,598 sequences dataset was used to calculate a dN/dS for *phoR* and other genes to increment the robustness of the measures and the number of variants per gene.

#### Transmission dataset

A dataset including samples taken over a 15-year period in the district of Malawi[39]. This dataset is enriched in transmission clusters and was used for the *phoR* transmission analysis.

#### MTBC most likely ancestral genome

The MTBC ancestor was derived in a previous publication by maximum parsimony and likelihood methods[33]. This ancestor is H37Rv-like in terms of genome structural variants, but H37Rv alleles were replaced by those present in the inferred common ancestor of all MTBC lineages.

### FASTQ mapping and variant calling for the MTBC strains

Fastq files were trimmed to remove low-quality reads using prinseq[57] and aligned to the MTBC most likely ancestral genome[33] using BWA-mem algorithm[58]. Alignments with less than 20x mean coverage per base were filtered out. The variant calling was performed using samtools[59] and VarScan[60]. Due to the low variability found in *M. tuberculosis*, to avoid mapping errors and false SNPs a variant was filtered out if: i) it was supported by less than 20 reads; ii) it was found in a frequency of less than 0.9; iii) it was found near indel areas (10 bp window); or iv) it was found in areas of high accumulation of variants (more than 3 variants in a 10 bp defined window). Variants were annotated using SnpEff[61]. Variants present in PE/PPE genes, phages or repeated sequences were also filtered-out, as they tend to accumulate SNPs due to mapping errors. High quality variant calls were combined in a non-redundant variant list and used to retrieve the most likely allele at each strain to generate a variant alignment.

### Phylogenetic inference and parsimony mapping of SNPs

In the MTBC dataset we identified 140,239 variants following the steps defined above. As we wanted to identify nucleotide variants due to recombination events, a stricter filtering was applied to remove putative recombination signal due to polymorphisms introduced by other causes. Variants related to antibiotic resistance were obtained from PhyResSe[62] and were removed from the analysis. Also, non-biallelic variants were removed from the analysis. To avoid false positives, we also removed positions in which a variant was called in at least one strain but also with a gap in at least another strain. To identify variants resulting from mapping errors we generated fragments of 50 bp downstream, upstream and midstream of the variant positions in the reference genome. With these fragments, we performed a BLAST search over the reference genome to check whether they mapped to other regions. Variants identified in reads that mapped to more than one region of the reference genome (query coverage per HSP over 98% and percentage of identical matches between the query and the reference genome of 98%) were removed from the analysis.

The remaining variants (94,780) were used to infer a phylogenetic tree using RAxML[63] with the GTRCATI (GTR + optimization of substitution rates + optimization of site-specific evolutionary rates) model of evolution and represented with the iTOL software[64]. Variants were mapped to the phylogeny using the Mesquite suite[65]. Homoplastic variants were identified based on parsimony criteria. Using these homoplastic variant positions, we looked for homoplastic runs. A homoplastic run was defined by two (or more) homoplastic variants found in the genome in correlative positions or with at least one variant between them. Variants present in the same homoplastic run were mapped on the phylogeny using Mesquite to look for coincident phylogenetic patterns.

### Linkage-disequilibrium calculation

Using the filtered variant positions (94,780), we used the PLINK software[66] to calculate the linkage-disequilibrium statistics D’ and R^2^. To estimate these values, we took into account variants with a minimum frequency of 0.01 and used a sliding window of 10 Kb. The results obtained were processed with R[67]. To plot the D’ and R^2^ pattern by variant distance, we calculated average D’ and R^2^ values for 50 bp sliding windows.

### Multiple alignment of *M. canettii* and MTBC

Seven *M. canettii* draft genomes were aligned to each other and to the ancestor of MTBC using progressiveMauve[68]. The segmented alignment obtained in XMFA format was converted to a plain Fasta format using the MTBC ancestor as reordering reference with a custom Perl script. Positions with gaps in the reference sequence were removed from the final alignment, so the resulting aligned genomes had the same size than the reconstructed MTBC ancestor (4,411,532 Mb). The MTBC pseudogenomes reconstructed from mapping to the MTBC ancestor from the different datasets described above were concatenated to the *M. canettii* alignment obtained in the previous step for further analyses.

From these alignments, homoplastic variants were identified using both, parsimony and maximum-likelihood approaches[69]. Both approaches agreed in identifying the same homoplastic variants.

### Recombination analyses and phylogenetic evaluation

Besides SNPs, linkage-disequilibrium analysis and Gubbins, RDP4[70] was used to detect recombination signal in the MTBC dataset. To mark the regions reported by Gubbins as potentially recombinant we required at least three of the methods implemented in RDP4 to agree in showing a significant signal.

Recombination was evaluated in the alignment containing 219 strains from MTBC and 7 *M. canettii* and in the one containing the MTBC ancestor and 7 *M. canettii*. Firstly, repetitive regions (i.e. PPE/PGRS) were masked from both alignments and, secondly, recombination events were inferred using Gubbins[71], which identifies clusters of high SNP density as markers.

Gubbins identified 70 potential recombinant regions in the alignment containing the 7 *M. canettii* strains and the MTBC ancestor. Four of these regions were obviated because they fell in regions deleted in several *M. canettii* strains. One more region was removed from the analysis because it was extremely short (41 bp) and we did not obtain reliable results in the subsequent analyses.

For the remaining 65 fragments a phylogeny was calculated using RAxML[63] and applying the GTRCATI model. Also, a reference phylogeny was calculated with the same method using the complete genomes after subtracting these 65 regions. This reference phylogeny had the same topology as the one obtained from the complete genomes. To test for phylogenetic incongruence between the putative recombination fragments and the genome phylogeny, we applied the Shimodaira-Hasegawa and Expected Likelihood Weight tests implemented in TREE-PUZZLE[72].

### Dating analyses

To infer the age of the 65 recombinant fragments we first reasoned that most of the mutations found were contributed by recombination and not by mutation once the fragment had been integrated in the genome. Thus, before dating the fragments we first removed all the homoplastic variants with other MCAN strain found in the fragments. The final alignments for the 65 fragments consisted of only those variants accumulated after the recombination event. We then used the non-recombinant part of the genome to infer a substitution rate assuming two different dating scenarios published for the tMRCA[6,73]. We ran BEAST[74] for each fragment pre-specifying monophyletic groups and substitution rate based on the non-recombinant genome phylogenetic reconstruction. We used an uncorrelated log-normal distribution for the substitution rate in all cases and a skyline model for population size changes. We ran several chains of up to 10E6 generations sampling every 1E3 generations to ensure independent convergence of the parameters. Convergence was assessed using Tracer[75]. For both evolutionary scenarios, the results obtained were largely congruent and proportional to the age limit imposed for the MTBC ancestor. As there is controversy about the correct MTBC ancestor age, the results were transformed to relative ages for plotting the final results.

### Gene ontology enrichment analysis

Genes present in the recombinant regions between MCAN and the MTBC ancestor were annotated using SnpEff[61]. A Gene Set Enrichment (GSE) analysis was performed to look for enriched gene functions in these regions. The BiNGO tool[76] was used to study the enrichment in certain functional categories comparing the most abundant terms in the recombinant regions with those contained in the complete annotation. The tool uses a hypergeometric test (sampling without replacement) and the BH correction for multiple testing comparisons. The Cytoscape program[77] was used to visualize the results.

### divSNP analysis

From the *M. canettii* and MTBC ancestor alignment, we extracted those positions having one variant in all the *M.canettii* strains and another variant in the MTB ancestor. The divSNP frequency by nucleotide was calculated by dividing the total number of divSNPs (5,688) by the total number of bases in the alignment. Next, the expected abundance of divSNPs for each gene was calculated by multiplying the nucleotide divSNP frequency by the number of nucleotides in each gene. From the expected and observed divSNP abundances, we used a Poisson distribution to calculate the probability of having the observed divSNPs by chance for each gene. We selected genes having a pFDR <=0,01 using the q-value from Storey method[78].

Complete mycobacterial genomes for reference strains[79](Additional file 2: Table S9) were downloaded from RefSeq and GenBank. The orthologous genes were obtained from the amino acid sequences and using the Proteinortho tool[80]. A gene was considered as orthologous based on reciprocal best hits in BLAST. BLAST analysis required a minimum identity of 25%, a query coverage of 50% and a maximum e-value of 1E-05. The orthologous genes were aligned using Clustal-omega[81] and the phylogenies were constructed using RAxML and applying the PROTCATIAUTO model. The reference phylogeny was constructed using only the core genome (proteins having orthologous in all the mycobacterial genomes downloaded) with RAxML using the same options as above. The reference and alternative phylogenies calculated with the orthologous genes for the divSNPs enriched genes were manually inspected to check for congruence.

### dN/dS analysis

The dN/dS statistics were calculated using R[67]. The potential synonymous and non-synonymous substitution sites for each region were calculated using the SNAP tool[82,83]. The dN/dS ratio for each region was calculated as follows:

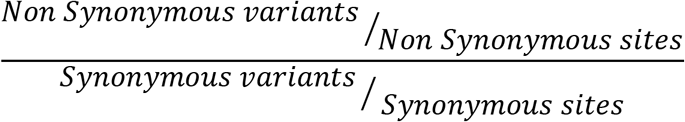

The dN/dS for the MTBC ancestor was calculated using the divSNPs while the dN/dS for the MTBC were calculated using 208,238 variants detected in coding regions from the 4,598 strains in the MTBC global dataset. To look for a robust comparison between both ratios, only genes having at least 1 synonymous and 1 non-synonymous variants were taken into account. To compare the dN/dS ratios, both were normalized by the genomic dN/dS for each taxon(0,24 for the MTBC ancestor and 0,59 for the MTBC). The difference between the dN/dS ratio was calculated by subtracting the MTBC dN/dS to that of the MTBC ancestor. The genes that account for the largest differences in the dN/dS were identified as outliers (Q2-1.5*IQR, Q3+1.5*IQR)[35] of the differences distribution.

### *phoR* positive selection analysis

Positive selection on *phoR* was tested using FUBAR[84] and BUSTED[85]. FUBAR was run with 5 MCMC chains of length 10,000,000. 1,000,000 states were used as burn-in and a Dirichlet prior of 0.5. BUSTED was run with default parameters.

To study the potential effect of *phoR* mutations on transmission efficacy we used the dataset from Guerra-Assunção *et al*.[39]. We identified SNPs in branches leading either to leaves or to transmission clusters. Transmission clusters were categorized in large, medium or small according to the number of isolates in the cluster (large = over 75^th^ percentile, medium=between 25^th^ and 75^th^ percentile, small = under 25^th^ percentile). Each gene was scored to check for accumulation of mutations in branches leading to large transmission clusters according to the expression:

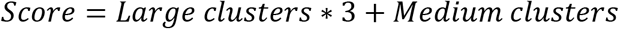

Genes with high mutation rates have a higher number of polymorphisms that could lead to a larger score by chance. To test the probability of obtaining the observed score by chance, a permutation test was carried out 10,000 times. Each of the identified SNP was randomly reassigned to the same branches and the score was recalculated for each gene. The expected score distribution for each gene was compared to the observed score to calculate the probability. This test was performed for transmission events defined at 10 SNPs.

The ages for the variant positions were calculated as node-to-tip distances. These distances were relativized to the maximum root-to-tip distance to obtain a relative age value in the [0 - 1] range. In order to have a common framework, a phylogeny was constructed including all the samples from the transmission and the reference datasets. The phylogeny was constructed using RAxML and applying the GTRCATI model. For each variant position we first identified the node in which the variant appeared. The node-to-tip distance was calculated afterwards for each node using the geiger package[86]. Distances were normalized to obtain a relative distance. Later, all the non-synonymous variants except the *phoR* polymorphisms were used as a reference set. The nonsynonymous *phoR* variants to be compared were categorized in two groups, those exclusive to the reference dataset[26] and those derived from the transmission dataset[39].

### PhoR domains structure representation

The PhoR domains structure was inferred by using PFAM(Finn et al. 2016) and SMART(Letunic and Bork 2018).

## DECLARATIONS

### DATA AVAILABILITY

The datasets used and/or analysed during the current study are available from the corresponding author on reasonable request.

## ACKNOWLEDGEMENTS

We thank Alberto Marina for advice in the interpretation of the PhoR molecular structure. This work was funded by projects the European Research Council (ERC) (638553-TB-ACCELERATE) and Ministerio de Economía y Competitividad (Spanish Government) research grant SAF2016-77346-R (to IC). BFU2014-58656-R and BFU2017-89594-R from Ministerio de Economía y Competitividad (Spanish Government) and PROMETEO/2016/122 from Generalitat Valenciana (to FGC). ACO is recipient of a FPU fellowship from Ministerio de Educación y Ciencia FPU13/00913 (Spanish Government). Swiss National Science Foundation (grants 310030_166687, IZRJZ3_164171, IZLSZ3_170834 and CRSII5_177163), the European Research Council (309540-EVODRTB) and SystemsX.ch (to SG).

## AUTHOR CONTRIBUTIONS

IC conceived this work. ACO, IC, LSB, SH, JC and FGC analyzed the data. ACO, IC, LSB, FGC wrote the first version of the draft. All authors critically reviewed and contributed to the final version of the manuscript.

